# Single-cell proteomics resolves major cell populations and protein-defined chondrocyte states in the adult human osteoarthritic joint

**DOI:** 10.64898/2026.09.25.754302

**Authors:** Leonie Müller, Jolet Y Mimpen, Caitlin JA Todd, Joseph Inns, Andrew M Frey, Sarah E Orr, Abby Brumwell, David J Deehan, Catharien MU Hilkens, Sarah JB Snelling, Matthias Trost, Sarah J Rice

## Abstract

**Objective:** Single-cell RNA sequencing has transformed our understanding of cellular heterogeneity within osteoarthritic (OA) joint tissues, yet transcript abundance provides only an indirect measure of cellular phenotype and function. Recent advances in mass spectrometry now enable proteins to be measured directly at single-cell resolution. Here, we applied single-cell proteomics (SCP) to primary human OA cartilage and synovium to determine whether cellular heterogeneity within the diseased joint can be resolved directly at the proteome level.

**Design:** Cartilage and synovium were obtained from patients undergoing total knee arthroplasty for OA. Low-input proteomic profiling was first performed to establish proteome depth from limited cell numbers and validate biologically relevant clustering. SCP was then applied to individual cells, followed by unsupervised clustering, differential protein abundance analysis and iterative biological annotation.

**Results:** Low-input proteomics identified 7,800 proteins from 1,000 cells while retaining tissue- and cell-associated biological differences. Across the SCP dataset, ∼2,300 unique proteins were detected, resolving chondrocyte, synovial fibroblast and immune-cell populations that closely paralleled those previously defined by single-cell transcriptomics. Higher-resolution analysis distinguished lining and sublining fibroblasts, discrete immune populations and multiple protein-defined chondrocyte states.

**Conclusions:** SCP enables direct characterisation of cellular heterogeneity within the human OA joint at the level of the proteome. By independently recovering established cellular populations while revealing protein-defined chondrocyte states, our findings establish SCP as a powerful complement to single-cell transcriptomics and open a new route to investigating functional cellular states and disease mechanisms that cannot be inferred from RNA abundance alone.

## Introduction

Osteoarthritis (OA) is a whole-joint disease in which pathological changes arise through the coordinated activity of multiple cell populations across cartilage, synovium, and other joint tissues. Single-cell resolution transcriptomic studies have transformed understanding of this cellular complexity. In articular cartilage, these approaches have identified distinct chondrocyte states associated with extracellular matrix production and remodelling, inflammation, cellular stress and hypertrophic differentiation[1]. Parallel studies of OA synovium have demonstrated substantial heterogeneity among both stromal and immune cells, including functionally distinct lining and sublining fibroblast populations[2]. Together, these studies establish cellular heterogeneity as a fundamental feature of OA joint tissues.

Despite such advances, transcript abundance provides only an indirect measure of cellular phenotype[3]. Protein abundance is shaped not only by transcription, but also by translational regulation, protein stability, degradation and post-translational modification. Consequently, RNA and protein abundance may diverge, particularly during dynamic cellular responses and transitions between disease-associated states. Proteins are also direct functional effectors of cellular behaviour and the targets of most therapeutic interventions. Therefore, analysis of protein abundance provides complementary biological information that cannot be inferred reliably from transcriptomic data alone.

Proteomics is routinely used to measure thousands of proteins, aiding understanding of cellular signalling cascades. However, until recently, proteomics analysis was restricted to bulk samples. Emerging single-cell proteomics (SCP) offers the potential to characterise cellular heterogeneity directly at the protein level[4]. Whereas transcriptomic workflows can profile tens of thousands of cells in parallel, mass-spectrometry-based SCP generally acquires one cell at a time. Further, unlike mRNA transcripts, proteins cannot be amplified before analysis. Each cell therefore constitutes a finite analytical sample, placing stringent demands on every step from sample preparation to computational analysis. Recent advances in low-input handling, trapped-ion mobility separation, parallel accumulation– serial fragmentation, and machine-learning-assisted peptide identification have substantially improved SCP depth and reproducibility [5].

Recent SCP studies have tracked how cellular protein content changes throughout the cell cycle[6], how cells react to inflammatory stimuli[7], how comparable proteome coverage is achieved across different source materials[8], and which protein signatures distinguish stem cells and early lineages[9]. In human skin tumours, this technology has already proven capable of resolving immune-cell populations, showing that SCP is translatable to clinical tissue biopsies[10]. However, it remains unknown whether contemporary SCP can recover the recognised cellular populations and molecular states of primary human OA tissues. Here, we evaluated whether contemporary SCP could resolve biologically meaningful cellular heterogeneity directly from primary human OA cartilage and synovium.

## Methods

Detailed methods are provided in the supplementary materials.

### Human tissue collection

Human knee articular cartilage and synovial tissue were obtained with informed consent from patients undergoing elective total knee replacement surgery for primary OA. Ethical approval was granted by the Newcastle and North Tyneside Research Ethics Committee (REC reference 19/LO/0389). Samples from seven donors (details in Table S1) were collected and processed to isolate donor-matched chondrocytes and synovial cells, as previously described[11,12].

### Cell isolation by FACS

Following tissue dissociation, synovial cells were FACS-separated into unlabelled synoviocytes, CD45^+^ immune cells and CD45^−^/PDPN^+^ fibroblasts, whilst chondrocytes were collected without further population selection. For low-input proteomics, 1,000 cells from each population were analysed from six tissue-matched donors. For SCP, CD45^+^ and CD45^−^/PDPN^+^ synoviocytes, and unlabelled chondrocytes from one additional donor were sorted individually into 96-well plates. For spectral library generation, we collected four 300-cell pools per population.

### Low-input proteomics

Samples were digested with trypsin/LysC and analysed by LC-MS/MS using an Evosep One coupled to a timsTOF HT with DIA-PASEF acquisition[5]. Raw data were analysed using DIA-NN[13]. Following quality filtering, protein intensities were log_2_-transformed and differential protein abundance and GO enrichment analyses performed as detailed in the Supplementary Methods.

### Single-cell proteomics

SCP was performed using cells from one biological donor. Individually sorted cells were digested in 96-well plates and analysed using the same LC-MS/MS platform. The parallel 300-cell samples were used to generate a spectral library in DIA-NN, which was subsequently used for analysis of the single-cell data. Contaminants and proteins without unique peptides were removed. Cells showing extreme deviations (>1.6 SD) in the number of proteins identified or total protein intensity were excluded as technical outliers.

### Cell clustering and annotation

Single-cell proteomic data were processed in R and RStudio using Seurat. Data were subjected to unsupervised clustering and UMAP dimensionality reduction. Broad cell identities and refined cellular states were assigned iteratively using protein abundance profiles, established musculoskeletal cell markers and published single-cell transcriptomic datasets. Full analytical and quality-control procedures are provided in the Supplementary Methods.

## Results

Low-input proteomics establishes proteome depth and a reference for single-cell analysis Before undertaking single-cell proteomics, we assessed proteome recovery from low input samples of 1000 unsorted chondrocytes or synovial cells, along with CD45-/PDPN^+^ synovial fibroblasts, or CD45^+^ synovial immune cells from each of six biological donors. Cartilage and synovium were enzymatically dissociated, FACS sorted and processed for LC–MS/MS analysis. A mean of 6822 proteins was identified in chondrocytes, 7311 in unlabelled synovial cells, 7115 in CD45−/PDPN+ synovial fibroblasts, and 6764 in CD45^+^ synovial immune cells (Fig. 1A). Protein identification numbers and intensity distributions were highly concordant between biological donors (Fig.S2A-C). Across the low-input dataset, 7862 proteins were identified, of which 6629 were detected in ≥75% of samples and 5021 in all samples (Fig. S2D, Table S2), demonstrating reproducible proteome coverage.

**Figure 1.**
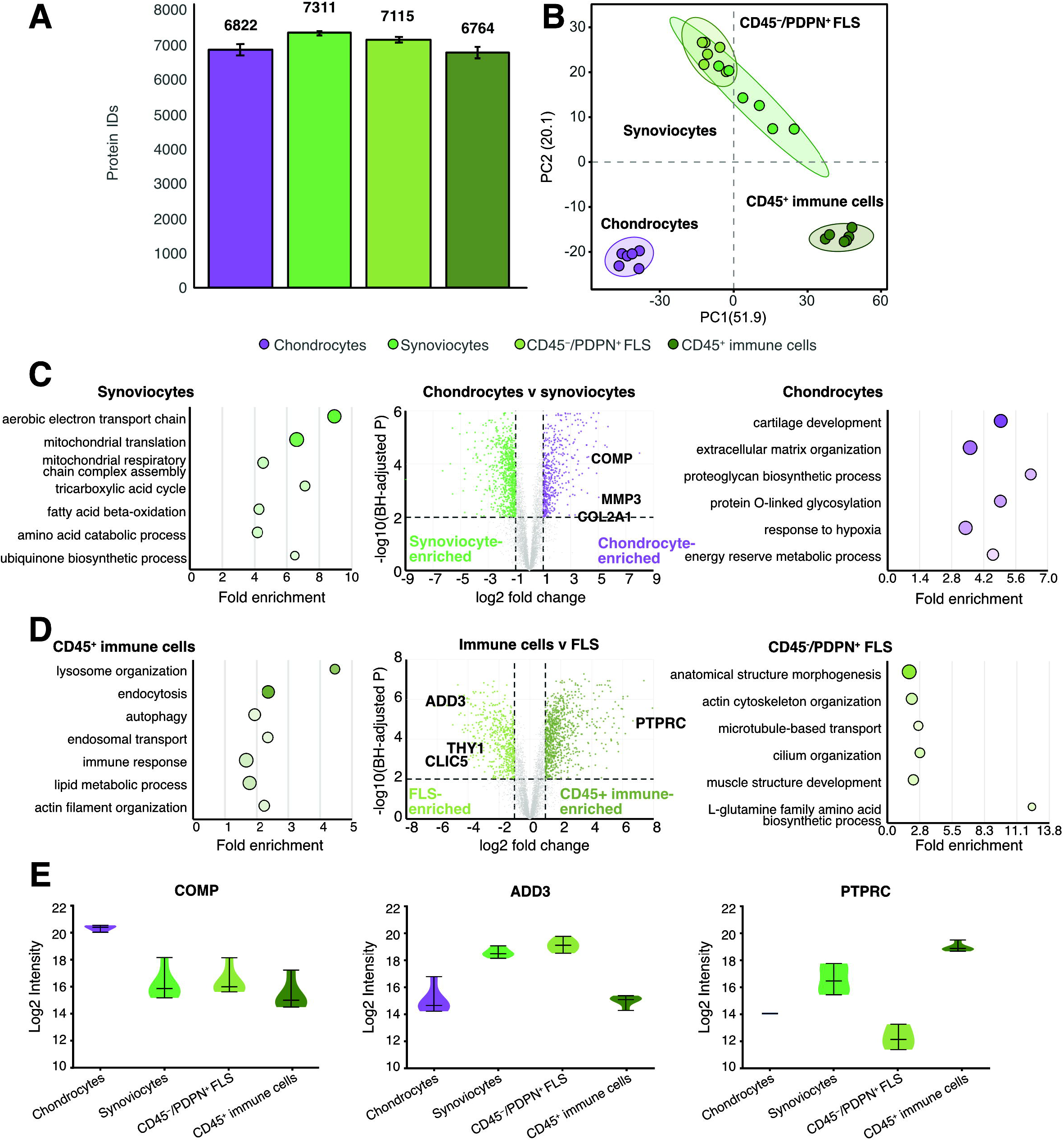
Low-input proteomic profiling establishes proteome depth and biological resolution in osteoarthritic joint cells. (A) Mean number of proteins identified from the peptide equivalent of 1,000 chondrocytes, synoviocytes, CD45^−^/PDPN^+^ synovial fibroblasts (FLS) and CD45^+^ immune cells from six tissue-matched donors. Error bars represent standard deviation. (B) Principal component analysis of low-input proteomic profiles demonstrating separation according to cell population across biological replicates. (C)Differential protein abundance analysis comparing chondrocytes and synoviocytes, together with representative significantly enriched GO Biological Process terms for proteins with higher abundance in each population. (D)Differential protein abundance analysis comparing CD45^+^ immune cells and CD45^−^/PDPN^+^ FLS, together with representative significantly enriched GO Biological Process terms for proteins with higher abundance in each population. For differential abundance analysis, dashed lines indicate BH-adjusted P <0.01 and |log_2_ fold change| >1. For GO enrichment plots, point size represents intersection size and colour represents −log10(FDR). (E)Protein abundances of COMP, ADD3 and PTPRC across the four low-input cell populations.

Principal component analysis (PCA) separated cartilage-derived chondrocytes, unlabelled synovial cells and the FACS-purified synovial populations, indicating that tissue- and cell-associated proteomic differences were retained at low cellular input (Fig. 1B). Unlabelled synovial cells occupied an intermediate position between the CD45^−^/PDPN^+^ fibroblast and CD45^+^ immune populations, consistent with their heterogeneous cellular composition.

Differential abundance analysis further demonstrated marked proteomic differences between cell populations (Fig. 1C–D; Table S3). Comparison of chondrocytes and synoviocytes identified 1,307 differentially abundant proteins (BH-adjusted *P* <0.01, |log_2_ fold change| >1), comprising 451 proteins with higher abundance in chondrocytes and 856 with higher abundance in synoviocytes. Comparison of CD45^+^ immune cells and CD45^−^/PDPN^+^ synovial fibroblasts identified 1,434 differentially abundant proteins, comprising 878 with higher abundance in immune cells and 556 with higher abundance in fibroblasts. Representative differences included higher COMP abundance in chondrocytes and higher ADD3 abundance in fibroblasts (Fig. 1C–E).

Gene ontology analysis of these directional protein sets identified distinct biological signatures (Fig. 1C–D; Table S4). Chondrocyte-enriched proteins were associated with cartilage development, extracellular matrix organisation (FDR=3.1x10^−7^) and proteoglycan biosynthesis, whereas synoviocyte-enriched proteins showed particularly strong enrichment of mitochondrial and metabolic processes, including aerobic electron transport (FDR=2.3x10^−54^). CD45+ immune-cell-enriched proteins were associated with lysosomal organisation (FDR=3.3x10^−16^), endocytosis, autophagy and endosomal transport, while fibroblast-enriched proteins included processes related to cytoskeletal organisation (FDR=7.3x10^−7^) and intracellular transport. Together, these findings demonstrate that low-input proteomics provided substantial and reproducible proteome depth while retaining biologically meaningful tissue- and cell-associated differences, providing a reference for subsequent single-cell analysis.

### Single-cell proteomics resolves the principal cellular populations of the OA joint

Next, we applied the SCP workflow to individual chondrocytes, CD45-/PDPN^+^ synovial fibroblasts, and CD45^+^ synovial immune cells, isolated from OA joint tissue of a single donor. Blank samples included as workflow controls showed significantly lower total protein intensities and protein numbers than the single-cell samples (Fig.S4A-B). Quality-control filtering based on the number of proteins identified and total protein intensity retained 78 chondrocytes, 39 CD45^−^/PDPN^+^ synovial fibroblasts and 36 CD45^+^ synovial immune cells for downstream analysis (Fig.S4C). Following filtering, an average of 900 proteins were identified in chondrocytes, 885 in CD45^-^/PDPN^+^ synovial fibroblasts, and 603 per CD45^+^ synovial immune cell (Fig. 2A, Fig.S4D, Table S5). As expected, the reduced proteome depth of single-cell measurements resulted in greater data missingness across cells compared with the low-input proteomic analysis (Fig.S4E-F). Despite this, unsupervised analysis readily resolved three major cellular compartments corresponding to chondrocytes, synovial fibroblasts, and synovial-derived immune cells (Fig. 2B).

**Figure 2.**
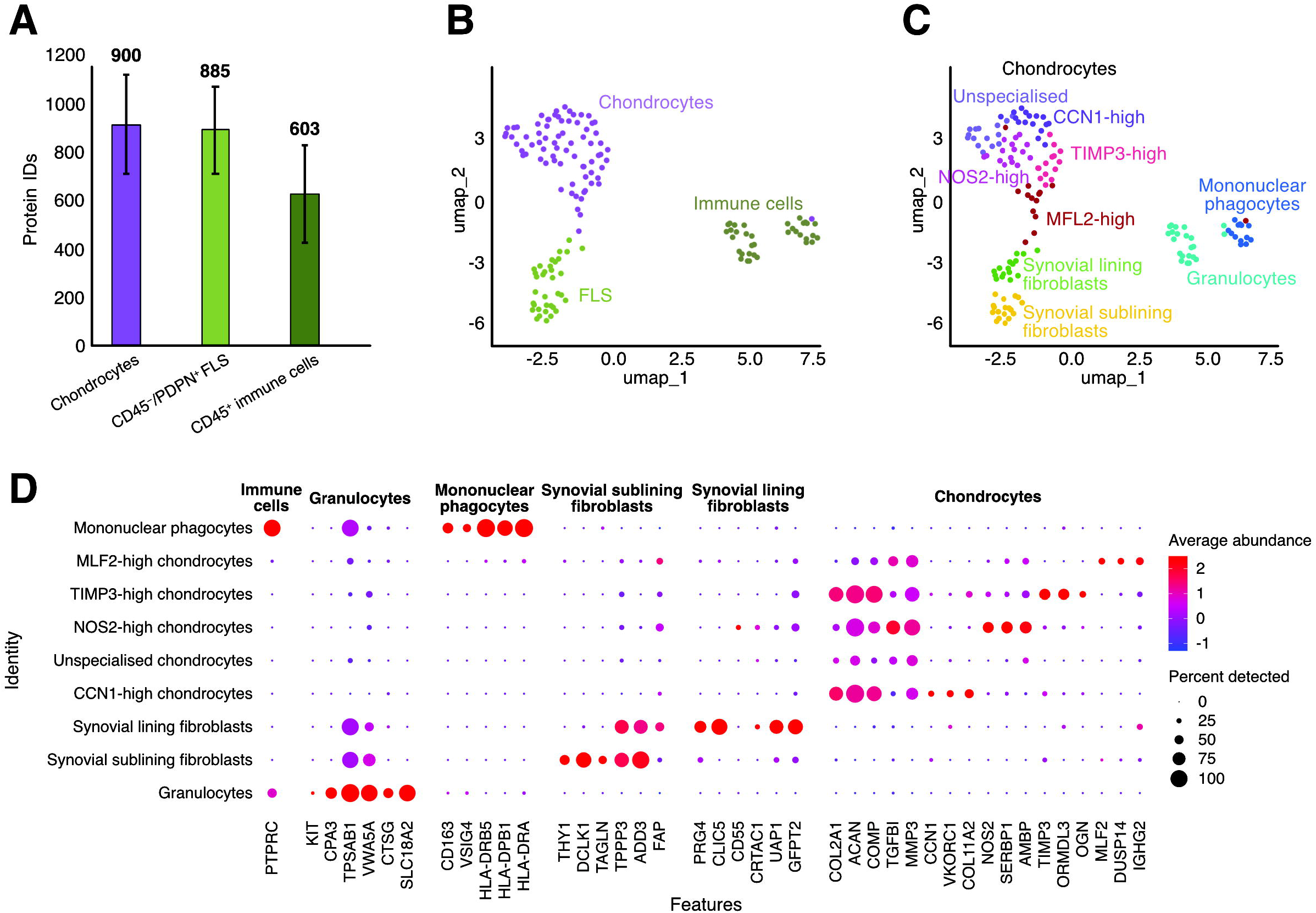
Single-cell proteomics resolves distinct cell populations in osteoarthritic joint tissues. (A)Mean number of proteins identified per cell in chondrocytes, CD45^−^/PDPN^+^ fibroblast-like synoviocytes (FLS) and CD45^+^ immune cells. Error bars represent standard deviation. (B)UMAP of single-cell proteomes showing separation of chondrocytes, FLS and immune cells. (C)UMAP showing higher-resolution protein-defined populations, including unspecialised, CCN1-high, NOS2-high, TIMP3-high and MLF2-high chondrocytes; synovial lining and sublining fibroblasts; granulocytes; and mononuclear phagocytes. (D) Dot plot showing selected marker proteins supporting cell-population annotation. Dot size represents the percentage of cells in which each protein was detected, and colour represents average scaled protein abundance.

Higher-resolution analysis demonstrated that the biological resolution extended beyond these broad cellular compartments. Synovial fibroblasts separated into PRG4-expressing lining and THY1-expressing sublining populations, closely paralleling the principal fibroblast populations previously identified by single-cell transcriptomic analysis of human OA synovium[2]. Within the immune compartment, the granulocyte population, likely consisting of mast cells, and mononuclear phagocyte populations were distinguished by characteristic protein abundance profiles, including TPSAB1 and major histocompatibility complex class II proteins, respectively (Fig. 2C–D).

### Single-cell proteomics resolves cellular heterogeneity in the OA joint

We next explored the heterogeneity within the chondrocyte compartment, which resolved into multiple cellular states characterised by distinct protein abundance profiles (Fig. 2C–D, Fig.S6, Table S6). These included CCN1-high and NOS2-high chondrocytes, together with populations characterised by elevated TIMP3 and MLF2 abundance. Pathway-enrichment analysis of the annotated single-cell populations is provided in Table S7. These populations were conservatively annotated according to their defining protein profiles rather than assigned to previously described transcriptomic chondrocyte states. Previous single-cell transcriptomic studies have similarly demonstrated substantial heterogeneity within human OA chondrocytes[1], although the protein-defined states identified here should not necessarily be expected to correspond directly to transcriptionally defined populations. Together, these findings demonstrate that cellular heterogeneity previously defined principally through single-cell transcriptomics can also be resolved directly from the proteome of individual primary OA cells.

## Discussion

The ability to measure proteins directly in individual cells provides an important complementary dimension to existing single-cell transcriptomic approaches in OA. Here, the recovery of recognised cellular compartments and synovial fibroblast populations previously identified by single-cell transcriptomic studies[1,2] demonstrates that biologically meaningful cellular organisation within primary human OA tissues can be resolved directly at the proteome level. Importantly, this was achieved using a label-free workflow, supporting the feasibility of applying SCP to clinically derived musculoskeletal tissues.

The value of SCP extends beyond reproducing transcriptionally defined cell populations. Protein abundance reflects the combined consequences of transcription, translation, degradation and protein stability, and therefore provides a distinct view of cellular phenotype that cannot necessarily be inferred from RNA abundance [3]. Although major cellular compartments and synovial fibroblast populations showed clear correspondence with those previously identified by single-cell transcriptomics, we deliberately did not assign the protein-defined chondrocyte states to existing transcriptomic classifications. Establishing the validity of clustering or equivalence between cellular types and states identified using different molecular modalities would require dedicated cross-modality integration and validation, particularly given the differences in feature number, analytical depth, and cellular scale between current SCP and scRNA-seq datasets. Moreover, one-to-one correspondence should not necessarily be expected, since proteomic states may capture cellular variation not represented by transcript abundance. The populations identified here are therefore described according to their defining protein profiles.

The importance of this additional molecular information in OA is illustrated by recent spatial mass spectrometry imaging of human osteochondral tissue, which identified spatially restricted extracellular matrix remodelling and post-translationally modified collagen peptides associated with diseased subchondral bone[14]. As proteomic depth and acquisition throughput increases, SCP should enable investigation of disease-associated cellular states defined by protein abundance and turnover, signalling pathways, and ultimately post-translational modifications. Integration of single-cell transcriptomic and proteomic measurements may therefore distinguish regulatory programmes that are transcriptionally encoded from those emerging principally at the protein level.

This study represents a proof-of-concept and has important limitations. SCP was performed in cells from a single donor and at a scale insufficient to assess inter-individual variation or the relative abundance of cellular states across OA. Protein detectability also varies according to abundance, extraction, digestion, and ionisation efficiency, as well as the dynamic range of the mass spectrometer. Consequently, undetected proteins cannot necessarily be considered absent, and measured intensities represent relative rather than absolute abundance. These considerations are particularly relevant when comparing protein-defined populations with transcriptomic datasets. The relatively small number of cells analysed also limits the confidence with which finer cellular populations can be annotated, particularly where annotation is supported by a limited number of detected marker proteins. Higher-resolution cell states identified here should therefore be interpreted cautiously pending validation in larger, multi-donor datasets.

Nevertheless, rapid improvements in analytical sensitivity, proteome depth and throughput are substantially expanding the scale and biological resolution achievable by SCP[6,7]. Larger, multi-donor studies will enable assessment of variation between patients, joint sites, and disease states, as well as identification of less abundant cellular populations and states. In conclusion, our findings establish the feasibility of SCP for resolving cellular heterogeneity directly at the protein level in primary human OA tissues and demonstrate its potential as a powerful complement to single-cell transcriptomics for investigating OA pathogenesis.

## Acknowledgements

We thank Prof Gisela Orozco for sharing the flow cytometry protocol and providing advice on the synovial cell-sorting strategy. The authors gratefully acknowledge the patients who donated tissue for this study. We acknowledge the Newcastle University Flow Cytometry Core Facility (FCCF) for assistance with the generation of Flow Cytometry data.

## Data availability

The mass spectrometry proteomics data generated in this study have been deposited with the ProteomeXchange Consortium via the PRIDE partner repository under accession PXD084266. Processed protein abundance data and associated analysis outputs are provided in the Supplementary Tables.

## AI use disclosure

ChatGPT (OpenAI) was used to assist with language editing and condensation of selected sections of the manuscript to meet the word-count requirements of the Brief Report format. All final text was reviewed and approved by the authors.

## Author contributions

LM, MT, CMUH and SJR conceived and designed the study. LM performed the single-cell proteomics experiments, including cell isolation, sample preparation and mass spectrometry workflows, and undertook the initial processing and analysis of the single-cell proteomic data. CT and AB contributed to sample retrieval and preparation. JI and AMF contributed to the development of the analytical pipeline and data analysis. SEO contributed to data analysis. JYM contributed to clustering, iterative biological annotation and interpretation of the single-cell proteomic data. SJBS contributed to development and refinement of the analysis pipeline and interpretation of the data. DJD contributed to sample acquisition and interpretation of the data. CMUH contributed to data interpretation. MT developed and supervised the analytical and technical proteomics pipelines and contributed to data interpretation. SJR supervised the study and contributed to data interpretation and manuscript preparation. MT, CMUH and SJR acquired funding supporting the study. All authors critically reviewed and approved the final manuscript.

