## Supplemental Material for "Single-cell proteomics resolves major cell populations and protein-defined chondrocyte states in the adult human osteoarthritic joint"

### Supplementary Methods

#### Human tissue processing and preparation of cell suspensions

Human knee articular cartilage and synovial tissue were obtained from seven donors undergoing total knee arthroplasty for primary OA. Donor metadata and allocation to the low-input and single-cell proteomics experiments are provided in Supplementary Table S1.

For chondrocyte isolation, articular cartilage was dissected from the joint, washed in PBS containing 200 IU/mL penicillin, 200 µg/mL streptomycin and 40 IU/mL nystatin, and cut into small pieces. Tissue was digested with 0.1% pronase in complete medium comprising DMEM (high glucose) supplemented with 10% FBS, 2 mM L-glutamine, sodium pyruvate, 200 IU/mL penicillin, 200 µg/mL streptomycin and 40 IU/mL nystatin for 1 h at 37°C with rotation in a CO₂-free incubator. Samples were washed twice with complete medium and subsequently digested with 0.1% collagenase in complete medium overnight at 37°C with rotation. Cell suspensions were passed through a 100-µm cell strainer, centrifuged at 1,200 rpm for 5 min, washed with PBS and resuspended in PBS[1].

For isolation of synovial cells, synovial tissue was dissected into small pieces, washed with PBS and digested with 1 mg/mL trypsin in reduced medium comprising DMEM (high glucose) supplemented with 100 IU/mL penicillin and 100 µg/mL streptomycin for 45 min at 37°C with rotation in a CO₂-free incubator. The supernatant was discarded and the tissue subsequently digested with 2 mg/mL collagenase in reduced medium for 3h at 37°C with rotation. Cell suspensions were passed through a 100-µm cell strainer, centrifuged at 1,200 rpm for 5 min, washed, and resuspended PBS[2].

For cell fixation, chondrocyte and synovial cell suspensions were fixed with 4% PFA for 10 min at 4°C with rotation. Fixed cell suspensions were centrifuged at 1,200 rpm for 5 min, washed twice and resuspended in ice-cold FACS buffer (PBS containing 1% FBS, 1% BSA and 0.01% sodium azide). Chondrocyte and synovial cell suspensions were stored at 4°C overnight before FACS.

#### Fluorescence-activated cell sorting for single-cell isolation

Cell suspensions were briefly vortexed and passed three times through a 100-µm cell strainer before counting and resuspension in FACS buffer at 1 × 10⁶ cells per 100 µL. Human TruStain FcX™ (BioLegend, #422302; 5 µL per 1 × 10⁶ cells) was added and samples incubated for 10 min at 4°C. Cells were subsequently stained with 5 µL per 1 × 10⁶ cells of CD45-FITC (BioLegend, #304006), CD31-Alexa Fluor 700 (BioLegend, #303134) and PDPN-PE/Dazzle 594 (BioLegend, #337028) for 30 min at 4°C [3]. Samples were washed once with FACS buffer before sorting using BD FACSAria II or BD FACSFusion cell sorters operated by the Newcastle University Flow Cytometry Core Facility. Single-stained compensation beads were used to generate fluorescence compensation controls and correct for spectral spillover.

The gating strategy is shown in Supplementary Figure S1. Cells were sequentially gated to exclude debris and doublets. Chondrocytes and unlabelled synovial cells were collected from the respective singlet populations. Synovial cells were additionally separated into CD45⁺ immune cells and CD45⁻/PDPN⁺ synovial fibroblasts.

For low-input proteomics, 2,500 chondrocytes and unlabelled synovial cells and 1,000 CD45⁺ synovial immune cells and CD45⁻/PDPN⁺ synovial fibroblasts were collected into protein low-bind tubes and stored at −80°C until processing. For SCP, individual chondrocytes, CD45⁺ synovial immune cells and CD45⁻/PDPN⁺ synovial fibroblasts were sorted into prepared 96-well plates containing hexadecane and proteomics digestion buffer. Four 300-cell pools of each population were collected in parallel for spectral-library generation.

**Low-input proteomics sample preparation and LC-MS/MS method**

Samples were thawed and centrifuged at 5,000 × *g* for 5 min. Proteins were digested in 0.2% DDM, 100 mM TEAB, 5ng/µL trypsin and 5 ng/µL LysC using 5 µL digestion buffer for 1,000-cell samples and 12.5 µL for 2,500-cell samples. Samples were incubated for 2 h at 45°C before cooling on ice and quenching with 0.1% formic acid (FA) and 1% DMSO. The peptide equivalent of 1,000 cells from each sample was loaded onto Evotips prepared according to the manufacturer's instructions. Thus, one 1,000-cell-equivalent sample from each of the four populations was analysed for each of six tissue-matched donors.

LC separation was performed using an Evosep One equipped with an Aurora Elite C18 column with integrated CaptiveSpray emitter (15 cm × 75 µm, 1.7 µm C18; IonOpticks, #AUR3-15075C18-CSI) maintained at 50°C. Samples were separated using the predefined Whisper Zoom 20 samples-per-day (SPD) method (58 min gradient). Mass spectrometry was performed using a timsTOF HT operated at a capillary voltage of 1,600 V, dry-gas flow of 3.0 L/min and dry temperature of 180°C. Data were acquired using a custom pyDIA-PASEF method comprising 16 variable ion-mobility and mass-to-charge (*m/z*) windows with two quadrupole positions per window. General MS settings were *m/z* 100–1,700 in positive-ion mode, with a TIMS ion-mobility range of 0.6–1.45 1/K₀ and ramp and accumulation times of 100 ms. MS/MS acquisition used an *m/z* range of 300–1,200, ion-mobility range of 0.6–1.4 1/K₀, cycle time of 1.2 s and collision energies of 20–59 eV. In-batch calibration was enabled.

#### Single-cell proteomics sample preparation and LC-MS/MS method

Single-cell proteomics was performed using the workflow described by Inns et al.[4]. Briefly, 96-well PCR plates were prepared by solidifying 5 µL hexadecane at the bottom of each well and overlaying this with 1 µL proteomics digestion buffer. Individual cells were FACS-sorted into the prepared plates at 4°C, followed by brief centrifugation. Samples were incubated for 2 h at 45°C before addition of 30 µL 0.1% FA/1% DMSO to terminate digestion. Plates were cooled on ice to re-solidify the hexadecane, inverted over prepared Evotips and centrifuged at 500 × *g* for 1 min at 4°C to transfer the peptide samples.

LC separation was performed using the Evosep One/Aurora Elite C18 configuration described above, using the predefined Whisper Zoom 40 samples-per-day method. Mass spectrometry was performed on the timsTOF HT using the same source conditions as for low-input proteomics. Single-cell data were acquired using a custom pyDIA-PASEF method comprising four variable ion-mobility and *m/z* windows with two quadrupole positions per window. General MS settings were *m/z* 100–1,700 in positive-ion mode, with a TIMS ion-mobility range of 0.6–1.45 1/K₀ and ramp and accumulation times of 200 ms. MS/MS acquisition used an *m/z* range of 300–1,200, ion-mobility range of 0.6–1.45 1/K₀, cycle time of 1.03 s and collision energies of 20–59 eV. In-batch calibration was enabled.

A total of 172 individual cells from one donor were analysed, comprising 88 chondrocytes, 42 CD45⁺ synovial immune cells and 42 CD45⁻/PDPN⁺ synovial fibroblasts. Four additional 300-cell pools were collected for each cell population for generation of the spectral library.

**Low-input proteomics data processing**

Raw MS data were analysed in DIA-NN v2.0.1, using the Swiss-Prot *Homo sapiens* reference proteome (UP000005640, including reviewed isoforms; downloaded September 2025), supplemented with the Hao Lab protein contaminant library. Searches were performed using trypsin/P specificity with one missed cleavage permitted, a maximum of one variable modification, peptide lengths of 7–30 amino acids and precursor charge states of 2–4. Modifications included N-terminal methionine excision, cysteine carbamidomethylation, methionine oxidation and N-terminal acetylation. Peptide- and protein-level false discovery rates were set at 1%, with mass and MS1 accuracies of 15 ppm, match-between-runs enabled and QuantUMS quantification. DIA-NN output was processed in R v4.4.0. Contaminants were removed [5] and proteins supported by fewer than two unique peptides were excluded. Protein intensities were log₂-transformed before downstream analysis. Data quality and reproducibility were assessed using protein-identification numbers, protein-intensity distributions, pairwise Pearson correlations and PCA.

Proteins with valid measurements in at least 70% of samples within at least one comparison group were retained for differential abundance analysis. Statistical testing was performed using two-sample t-tests with Benjamini-Hochberg correction for multiple testing. Proteins with |log₂ fold change| >1 and adjusted *P* <0.01 were separated according to direction of effect and subjected to Gene Ontology Biological Process enrichment analysis using g:Profiler. Enrichment was performed using genes represented in the detected low-input proteome as the statistical background, with Benjamini-Hochberg false discovery rate correction for multiple testing. GO terms with FDR <0.05 were considered significantly enriched. Complete differential abundance and enrichment results are provided in Supplementary Tables S3 and S4, respectively. g:Profiler production API build e114_eg62_p19_27110d83 (Ensembl 114; GO classes release 23 January 2026) was used. Proteomics data are deposited with the ProteomeXchange Consortium via the PRIDE repository under accession PXD084266.

**Single-cell proteomics data processing**

Raw data from the 300-cell pools collected alongside the single cells were first searched library-free in DIA-NN[6]against the same reference database and using the same search parameters as the low-input proteomic dataset. The resulting spectral library was subsequently used for library-based searching of the single-cell data in DIA-NN. Search outputs were processed in R v4.4.0 and contaminant proteins removed.

To assess the balance between proteome depth and identification confidence, protein groups were independently filtered using minimum criteria of either one or two unique peptides per protein. Requiring two unique peptides reduced the number of retained proteins by 24%, with a comparable effect across all three cellular populations (Fig. S3A–B). PCA produced highly concordant biological separation using either filtering criterion (Fig. S3C–D), indicating that the principal cellular structure was robust to peptide-filtering stringency. Subsequent analyses therefore used proteins supported by at least one unique peptide to maximise proteome coverage in the single-cell dataset.

Single cells were subjected to quality-control filtering based on the number of proteins identified and total protein intensity within each FACS-defined population. Blank/control wells and cells showing extreme deviations (>1.6 SD from the population mean) in either metric were excluded from downstream analyses. Following QC, 153 of 172 cells were retained, comprising 78 chondrocytes, 39 CD45^−^/PDPN⁺ synovial fibroblasts and 36 CD45⁺ synovial immune cells (Fig. S4).

#### Dimensionality reduction, clustering and cluster optimization

Single-cell protein abundance data were analysed using Seurat v5.5.0[7] in R (v4.6.1) and RStudio Server (v2026.04.0). Data were log-normalised and 500 variable protein features selected using the variance-stabilising transformation (vst) method before scaling. PCA was performed using 50 principal components (PCs), with inspection of the elbow plot used to select the first 15 PCs for downstream analysis.

A nearest-neighbour graph was constructed and unsupervised clustering performed across resolutions from 0 to 4.0. Cluster stability across resolutions was assessed using clustree v0.5.1[8]. A final resolution of 2.5 was selected because it retained separation of the three major FACS-defined cellular compartments while resolving biologically interpretable subpopulations (Fig. S5). UMAP was used for two-dimensional visualisation.

Differentially abundant proteins between clusters were identified using Seurat's FindAllMarkers() function with a Wilcoxon rank-sum test. Analyses were restricted to proteins detected in at least 25% of cells within the population of interest and a minimum log₂ fold-change threshold of 0.25. Differential protein abundance was visualised using dot plots, heatmaps, and FeaturePlots (Figs S6–S7; Table S6).

Broad cellular identities were initially assigned using detected canonical protein markers and were confirmed using FACS origin. Cluster annotations were subsequently refined iteratively using differentially abundant proteins, protein-marker profiles, and comparison with previously characterised musculoskeletal cell populations and published single-cell transcriptomic datasets. Synovial fibroblast and synovial immune subpopulations were annotated according to established lineage-associated protein profiles, whereas the suggested chondrocyte states were conservatively named according to their defining protein abundance profiles rather than assigned directly to existing transcriptomic classifications.

Pathway enrichment was performed using gprofiler2 v0.2.4 [9] using the top 100 proteins ranked by average log₂ fold change, restricted to proteins with an average log₂ fold change ≥0.5. Multiple-testing correction was performed using the FDR method, and complete enrichment results are provided in Supplementary Table S7.

### Supplementary Figures

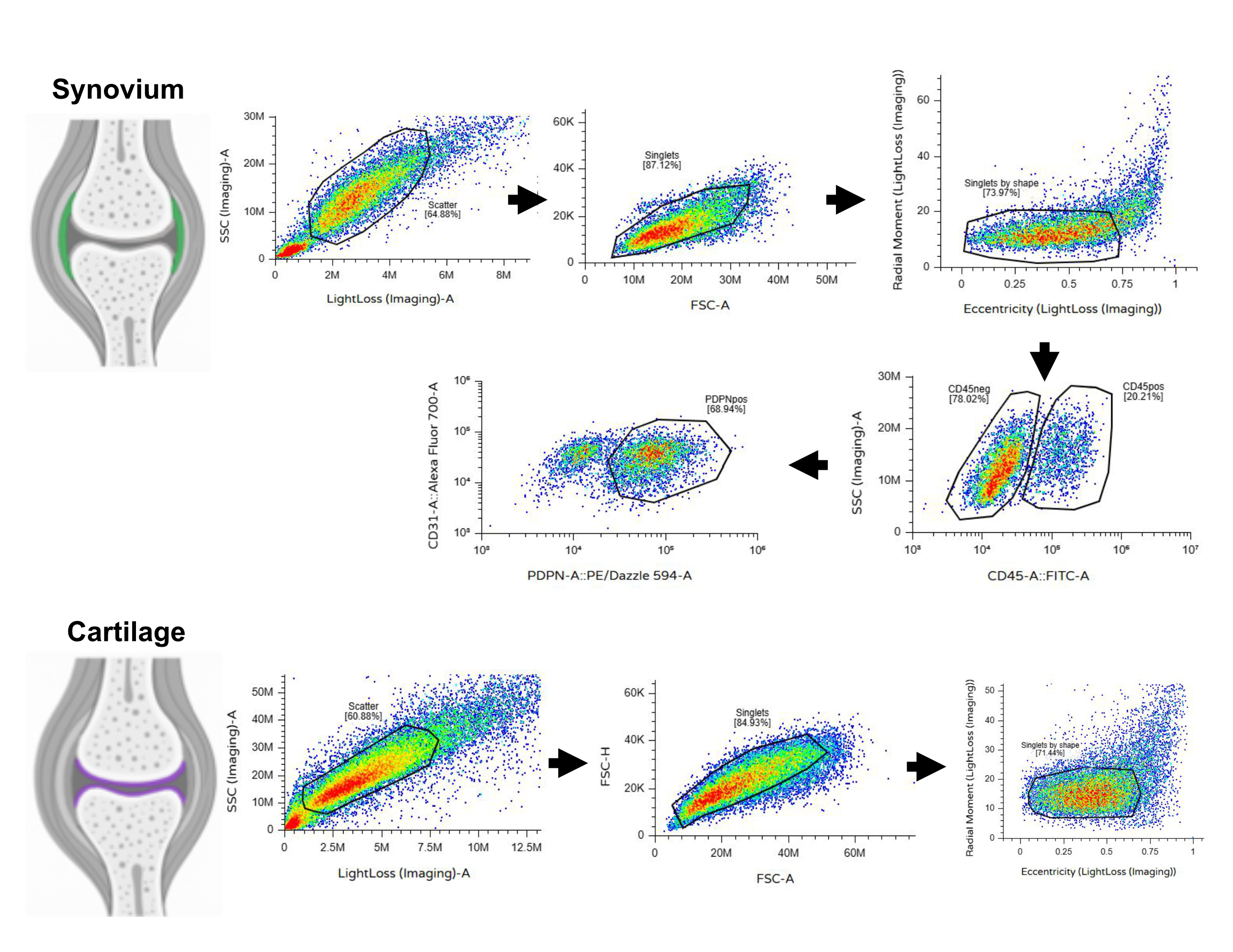

#### Supplementary Figure 1. FACS gating strategy for isolation of chondrocytes and synovial cell populations

Representative FACS gating strategy for chondrocytes and synovial cells. Samples were sequentially gated based on scatter characteristics to exclude debris and identify singlets. Chondrocytes were collected following singlet gating. Unsorted synovial cells were also collected following singlet preparation, but further gating for sub-populations was applied to obtain CD45⁺ immune cells and CD45⁻/PDPN⁺ synovial fibroblasts.

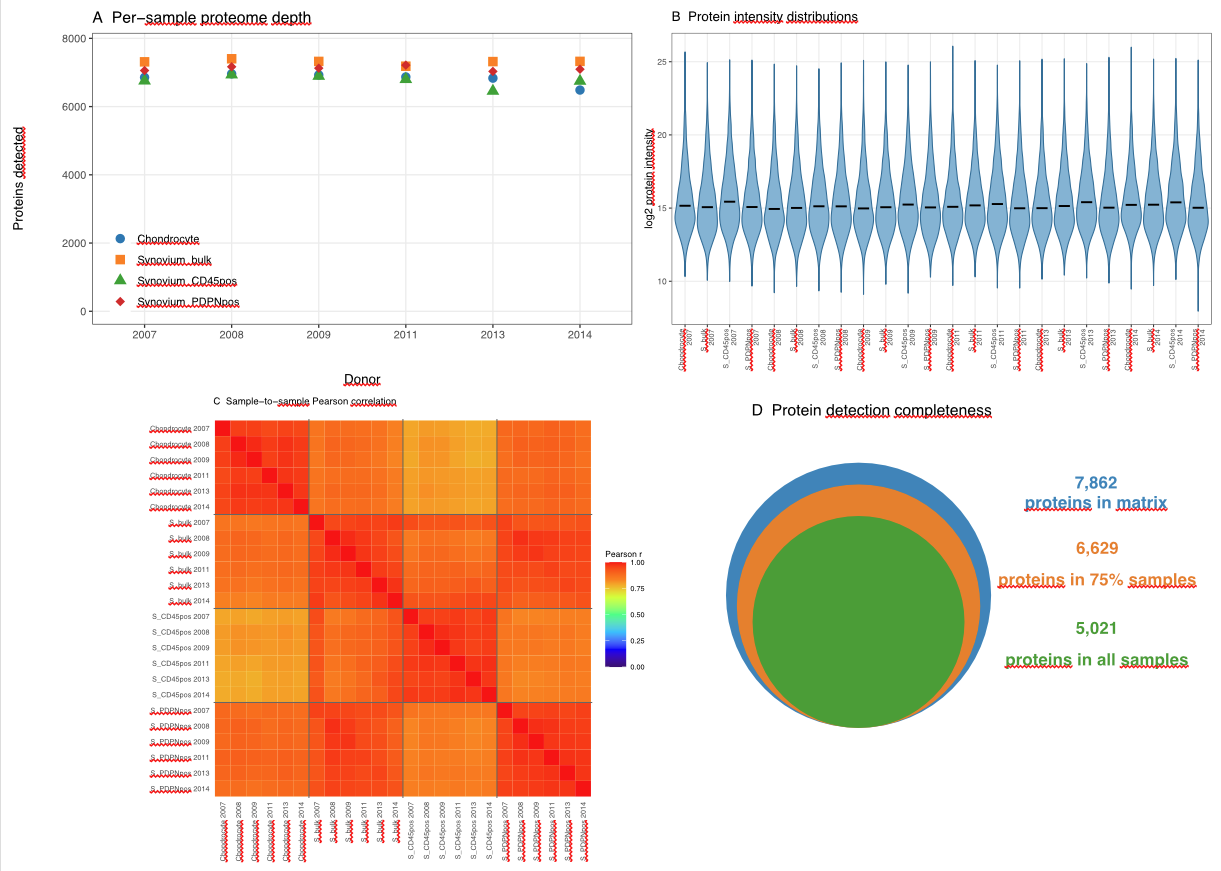
 **Supplementary Figure 2. Quality-control overview of the low-input proteomics dataset.**

(A) Number of proteins identified per sample, grouped by donor (n = 6) and distinguished by cell population. (B) Distribution of log₂-transformed protein intensities across the 24 individual samples. (C) Pairwise Pearson correlation coefficients of log₂-transformed protein intensities across all 24 samples. (D) Protein identification completeness across the low-input dataset. Nested areas represent the 7,862 protein groups identified across the experiment, of which 6,629 were identified in ≥75% of samples and 5,021 were identified in all 24 samples.

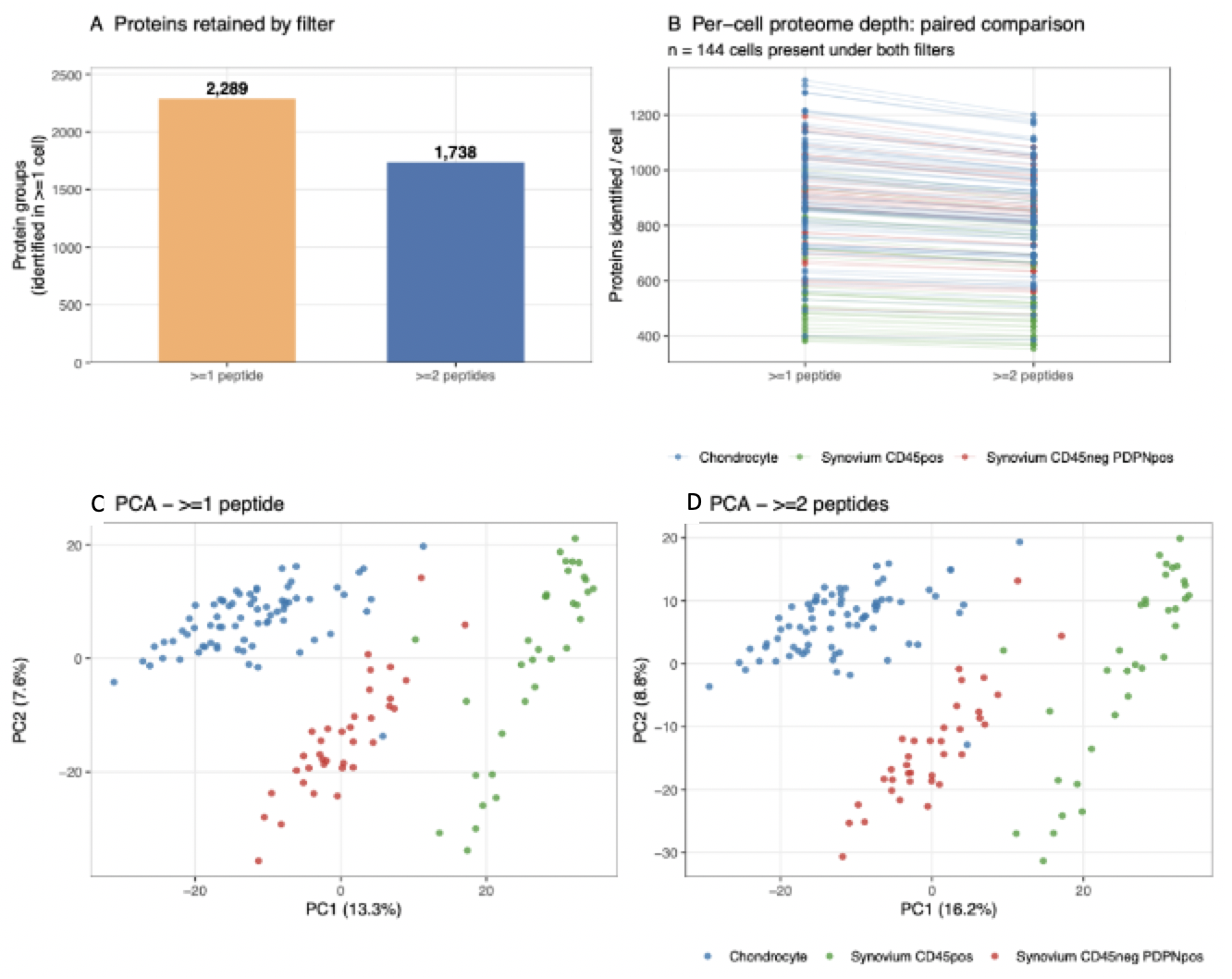
 **Supplementary Figure 3. Effect of one- versus two-peptide protein identification criteria.**

**(A)** Total number of protein groups retained using ≥1- and ≥2-unique-peptide identification criteria. **(B)** Number of proteins identified per cell under each filtering criterion; lines connect measurements from the same cell. **(C–D)** Principal component analysis (PCA) of single-cell proteomic profiles based on scaled protein intensities following filtering using the ≥1-unique-peptide **(C)** and ≥2-unique-peptide **(D)** criteria.

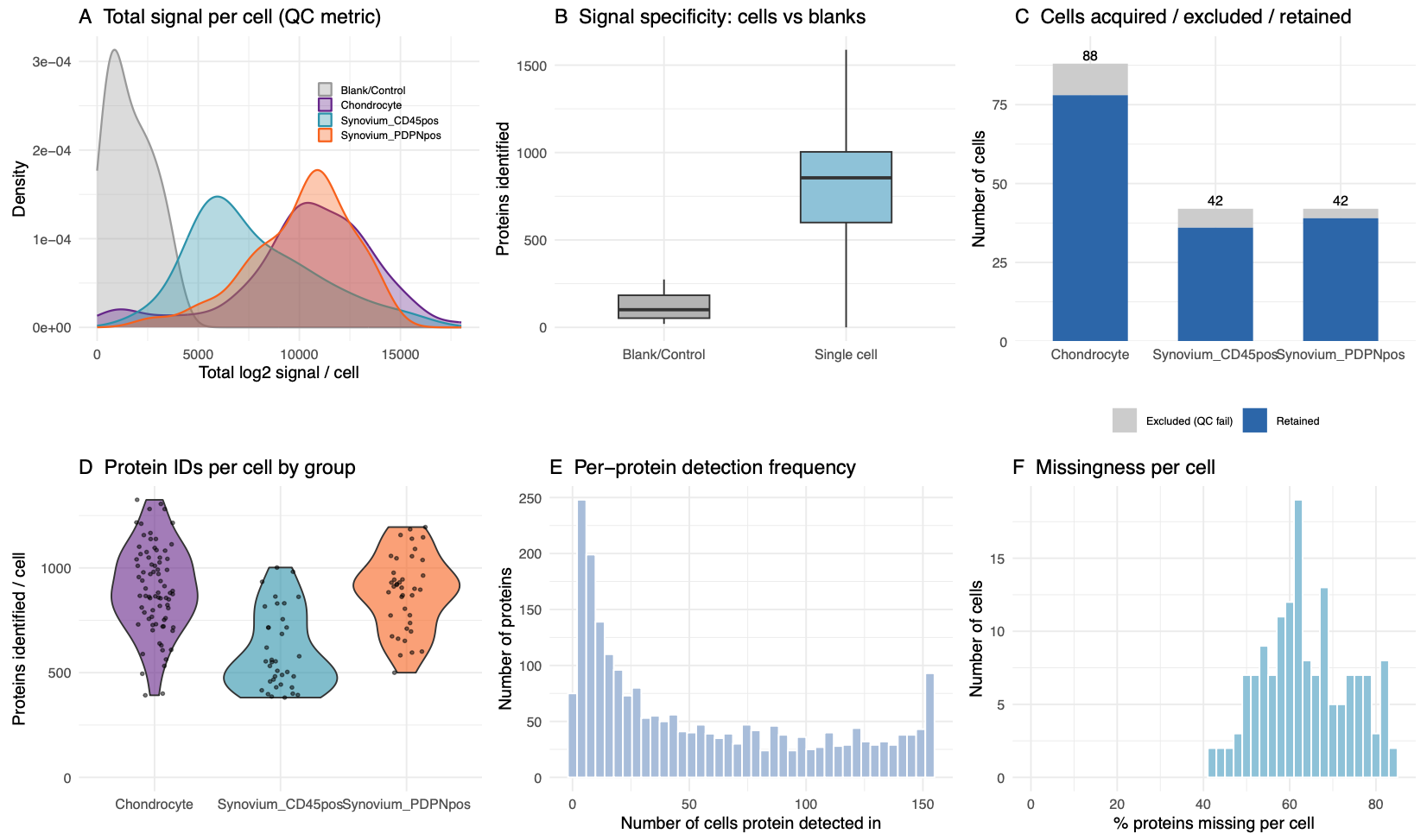

##### Supplementary Figure 4. Quality-control overview of the single-cell proteomics dataset.

**(A)** Distribution of total protein intensity across single-cell and blank/control wells. **(B)** Number of proteins identified in single-cell and blank/control wells. **(C)** Number of single cells acquired (grey) and retained (blue) following quality-control filtering. **(D)** Number of proteins identified per cells that were retained after filtering. Cells were retained when both the number of proteins identified and total protein intensity fell within 1.6 standard deviations (SD) of the mean for their FACS-defined cell population. **(E)** Detection frequency of individual proteins across all QC-passing cells. **(F)** Distribution of protein missingness among QC-passing cells, expressed as a percentage of the total quantified single-cell proteome. In total, 172 single cells and eight blank/control wells were analysed; 153 cells passed QC filtering and were retained for downstream analysis.

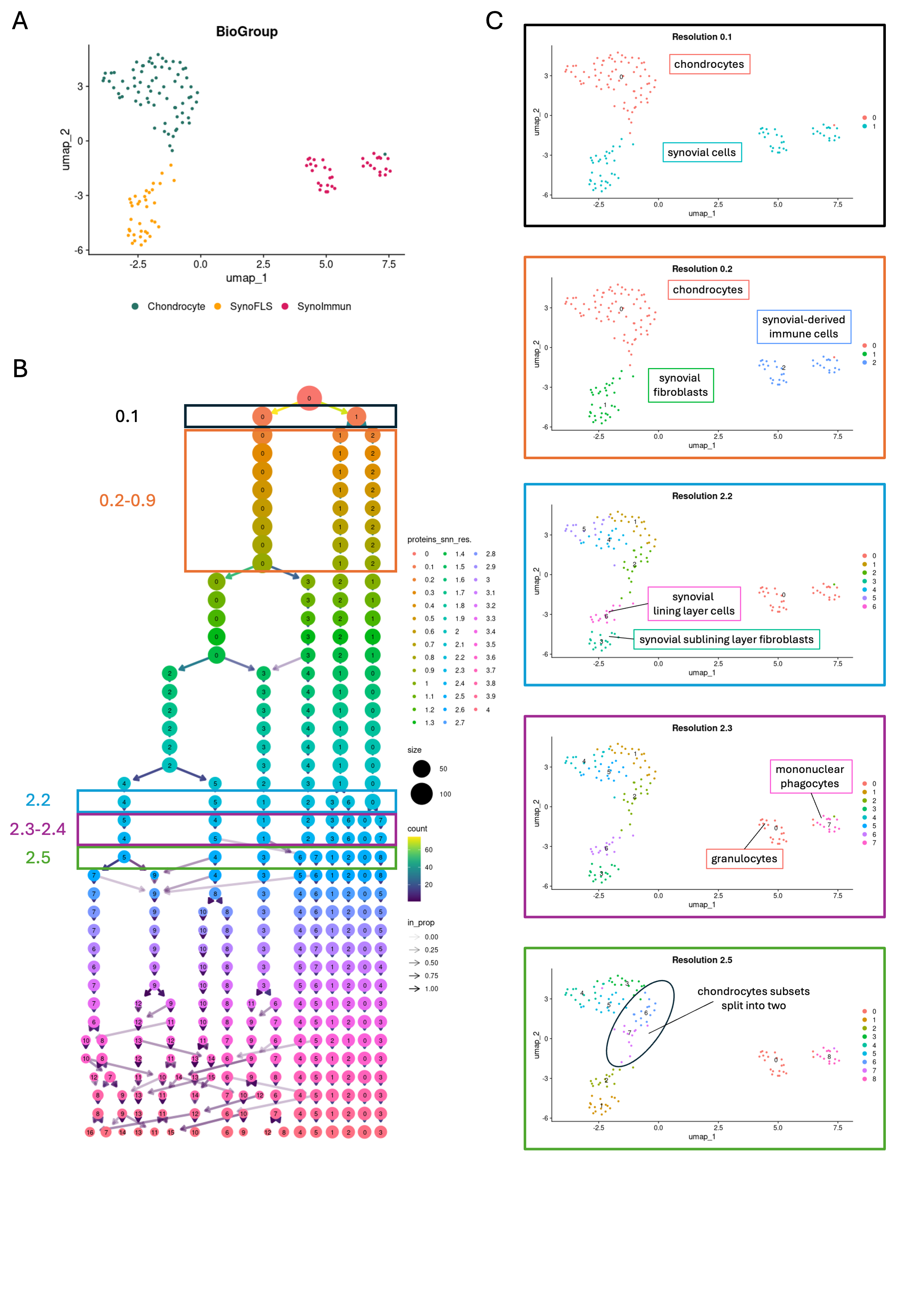

**Supplementary Figure 5. Clustering optimisation and selection of clustering resolution.**

(A) UMAP of single cells coloured according to their FACS-defined cell population of origin. (B) Clustree analysis showing the relationships between clusters across clustering resolutions from 0.0 to 4.0. (C) UMAPs illustrating the progressive resolution of cellular populations with increasing clustering resolution. At resolution 0.1, cells separated into cartilage-derived and synovial-derived populations. At resolutions 0.2–0.9, the three major FACS-defined populations corresponding to chondrocytes, synovial fibroblasts and synovial immune cells were resolved. At resolution 2.2, synovial fibroblasts separated into lining and sublining populations, while at resolutions 2.3–2.4, synovial immune cells separated into granulocyte and mononuclear phagocyte populations. Resolution 2.5 additionally resolved distinct chondrocyte states that remained stable across higher resolutions. Resolution 2.5 was therefore selected for downstream annotation as it provided biologically interpretable subdivision of the major cell populations, supported by differentially abundant protein profiles (Fig. S7), without further fragmentation at higher resolutions.

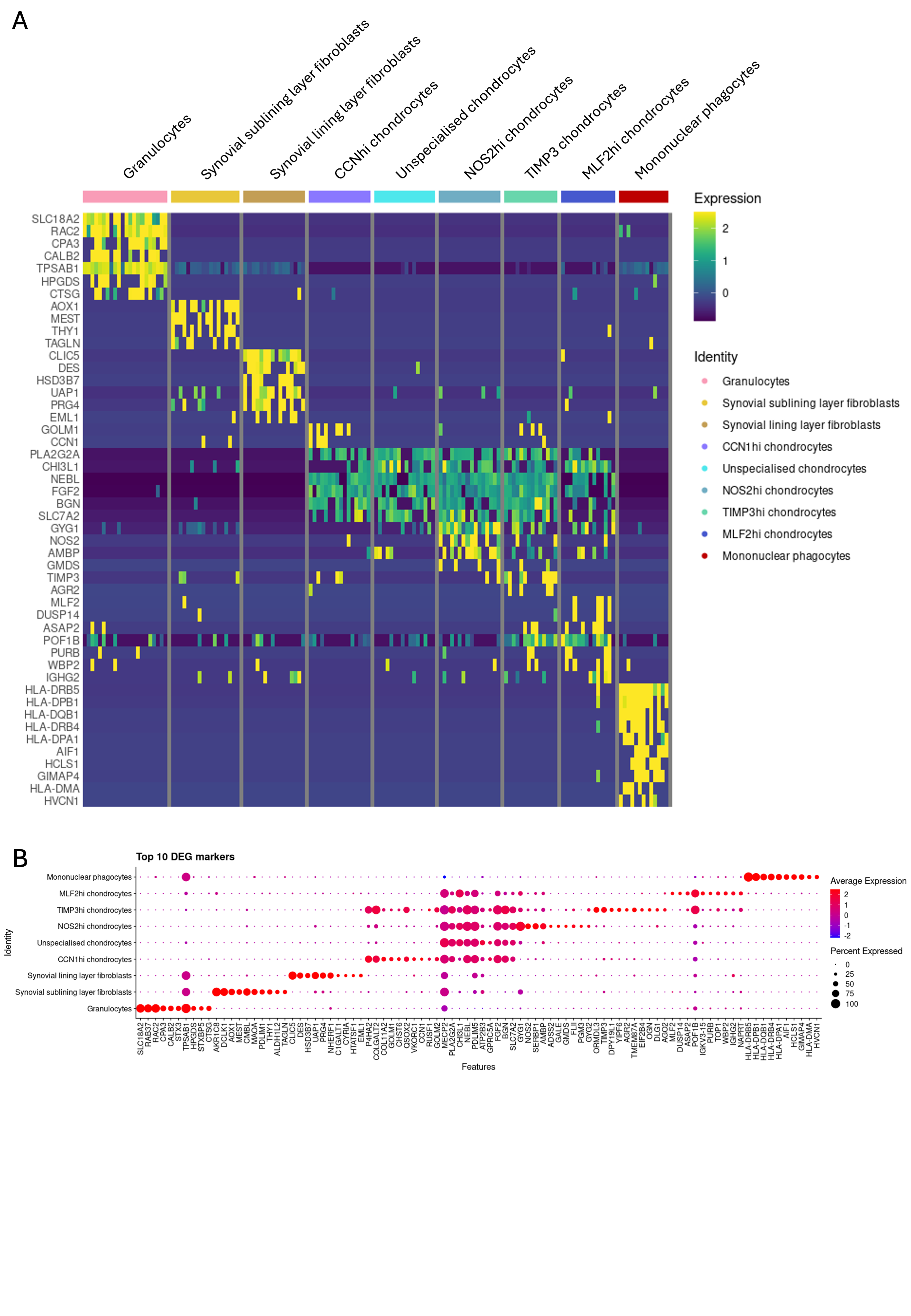

**Supplementary Figure 6. Differentially abundant proteins across single-cell clusters.**

**(A)** Heatmap showing the top 10 differentially abundant proteins for each cluster, identified by comparison of each cluster with all other cells. Protein abundance is scaled across cells for visualisation. **(B)** Dot plot showing the top 10 differentially abundant proteins for each cluster. Dot size represents the percentage of cells within each cluster in which the protein was detected, and colour represents average protein abundance.

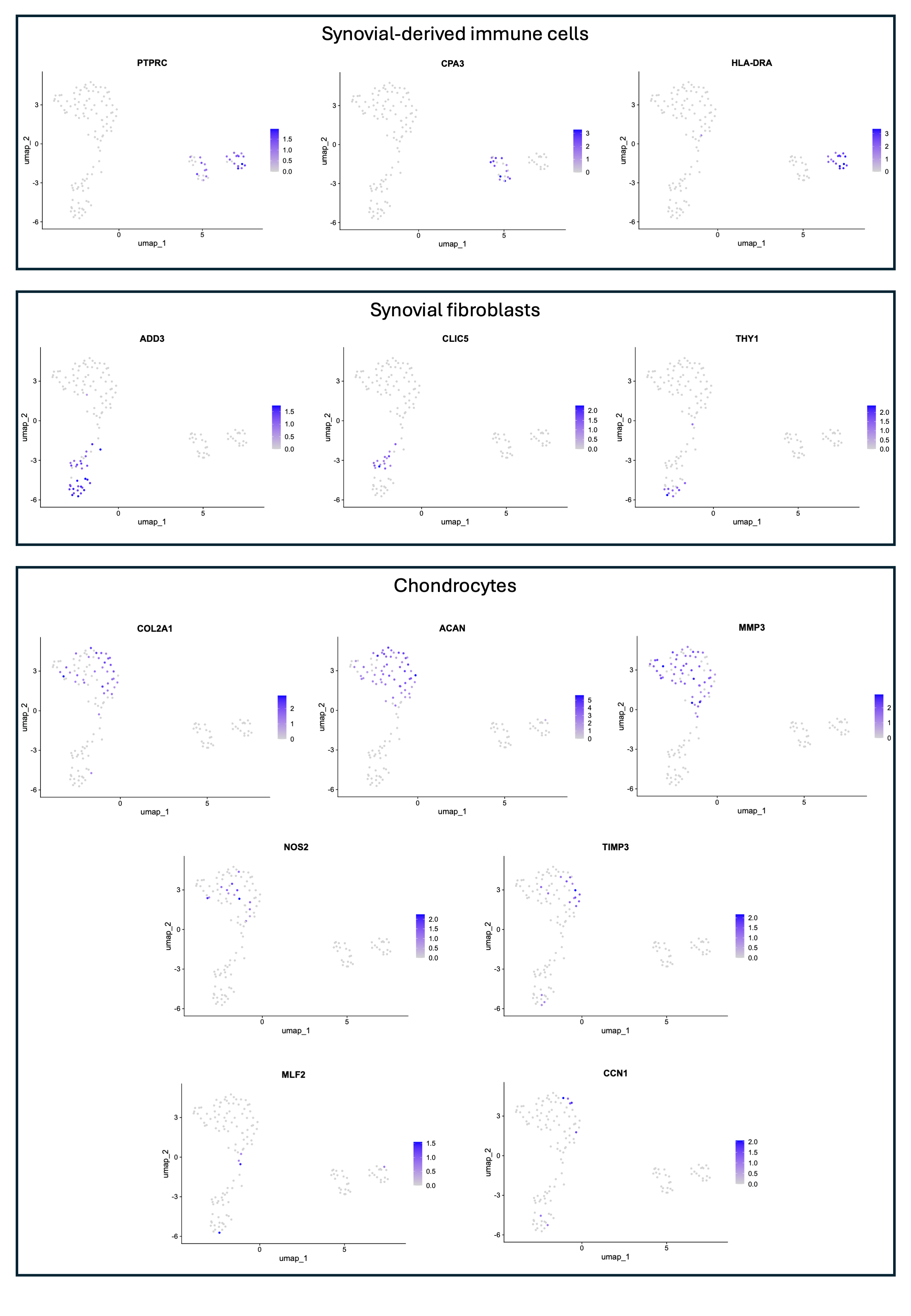

**Supplementary Figure 7. Protein-marker profiles supporting single-cell cluster annotation.**

Feature plots showing the abundance of selected proteins used to support annotation of single-cell clusters. Synovial-derived immune cells showed abundance of the pan-leukocyte marker PTPRC, with CPA3 enriched in the population annotated as granulocytes and HLA-DRA enriched in mononuclear phagocytes. Synovial fibroblasts showed ADD3 abundance, with CLIC5 enriched in lining fibroblasts and THY1 enriched in sublining fibroblasts. Chondrocytes showed abundance of cartilage-associated proteins COL2A1, ACAN and MMP3. Protein-defined chondrocyte states were further characterised by differential abundance of NOS2, TIMP3, MLF2 and CCN1.
